## Supplementary material for "RatXcan: A framework for cross-species integration of genome-wide association and gene expression data"

### Supplementary Figures

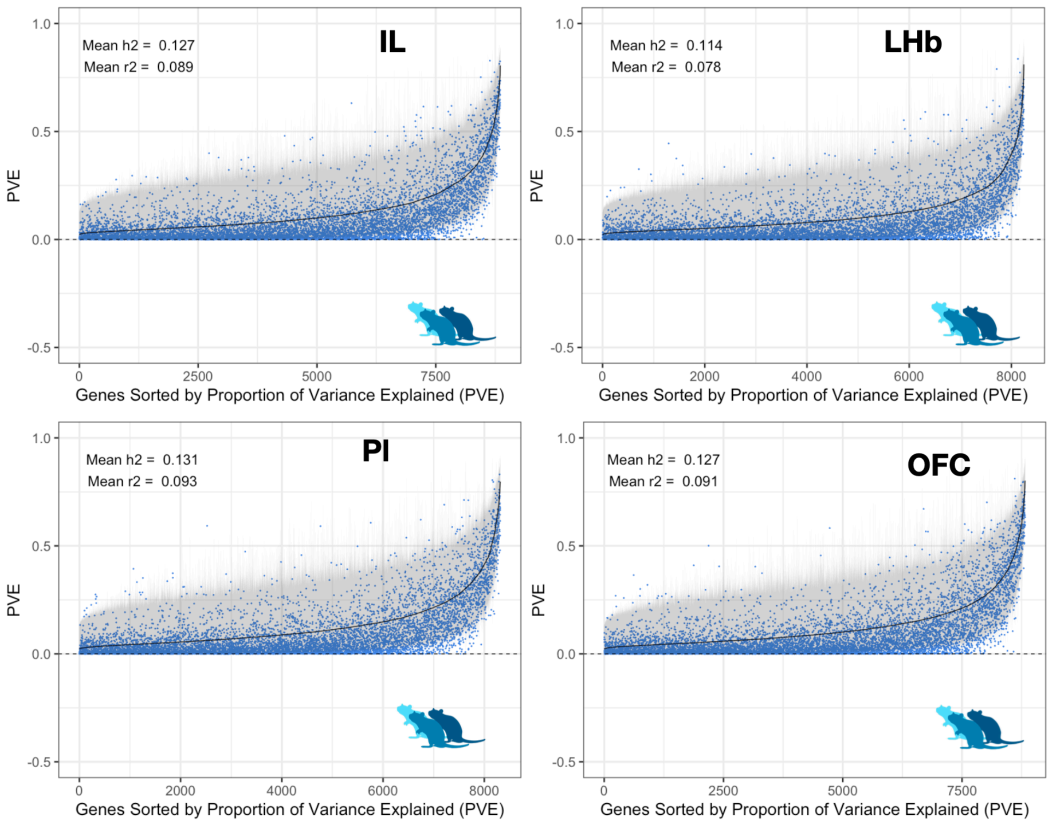


**Figure S1: Gene expression was heritable 8.86-10.12% and comparable across several brain tissues tested (Infralimbic Cortex, IL; Lateral Habenula, LHb; Prelimibic Cortex, PL; Orbitofrontal Cortex, OFC) in rats.** We refer to heritability ($h^2$, cis-heritability within 1Mb) as the proportion of variance explained (PVE). Across all brain tissues tested, heritability estimates were significantly correlated ($R$ = 0.58-0.83, $P < 2.20 \times 10^{-16}$).

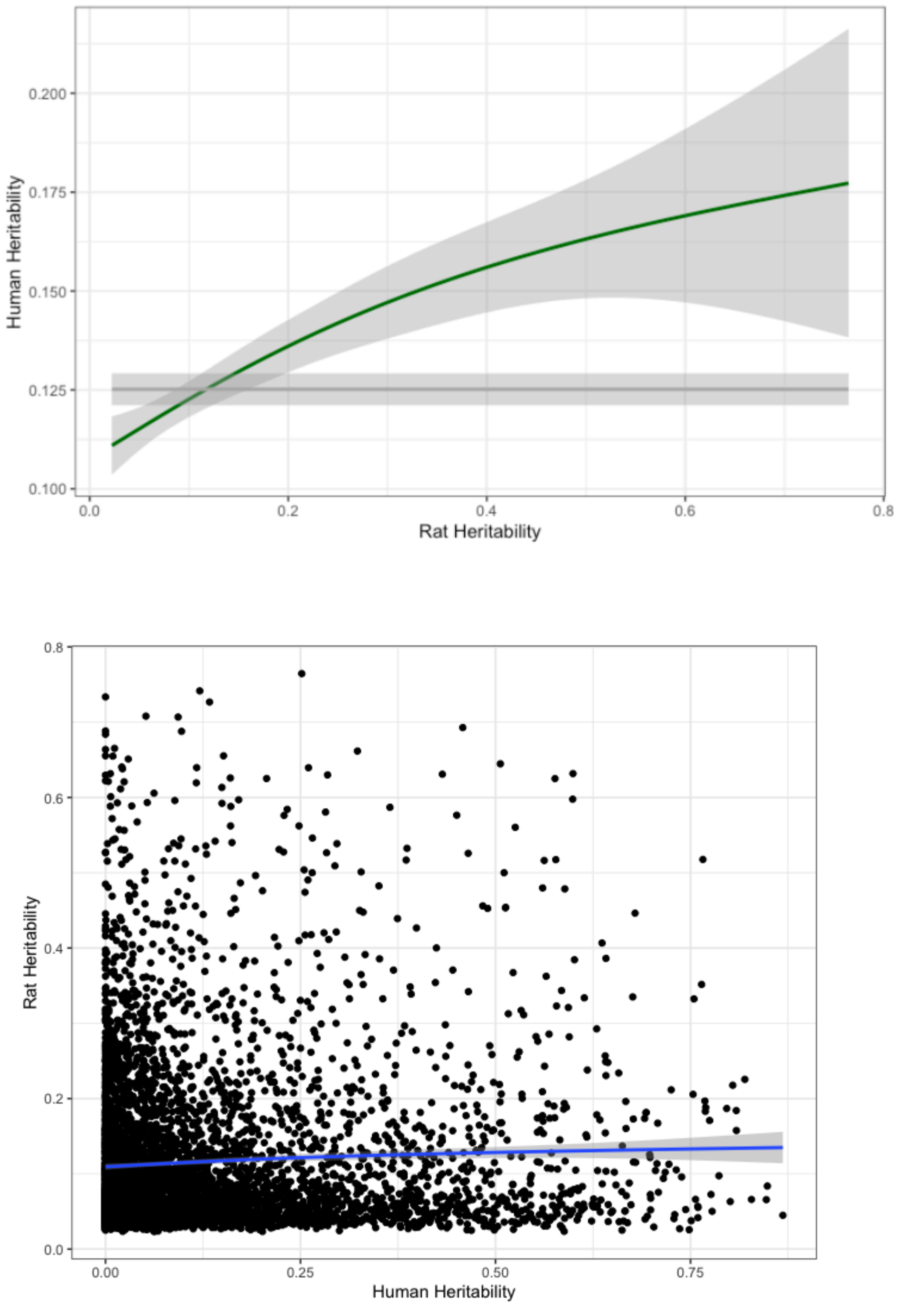


**Figure S2: Heritability of gene expression was correlated between rats and humans.** We found a significant correlation ($R$ = 0.07, $P$ = 4.34 x $10^{-12}$) between heritability estimates in rats and humans. Confidence intervals are represented as gray bars. The gray line represents the null distribution. Top panel shows the smoothed lines with loess (Local Polynomial Regression Fitting) implemented in the ggplot2 package in R. The bottom panel shows the same figure with all the points in addition to the smoothed curve.

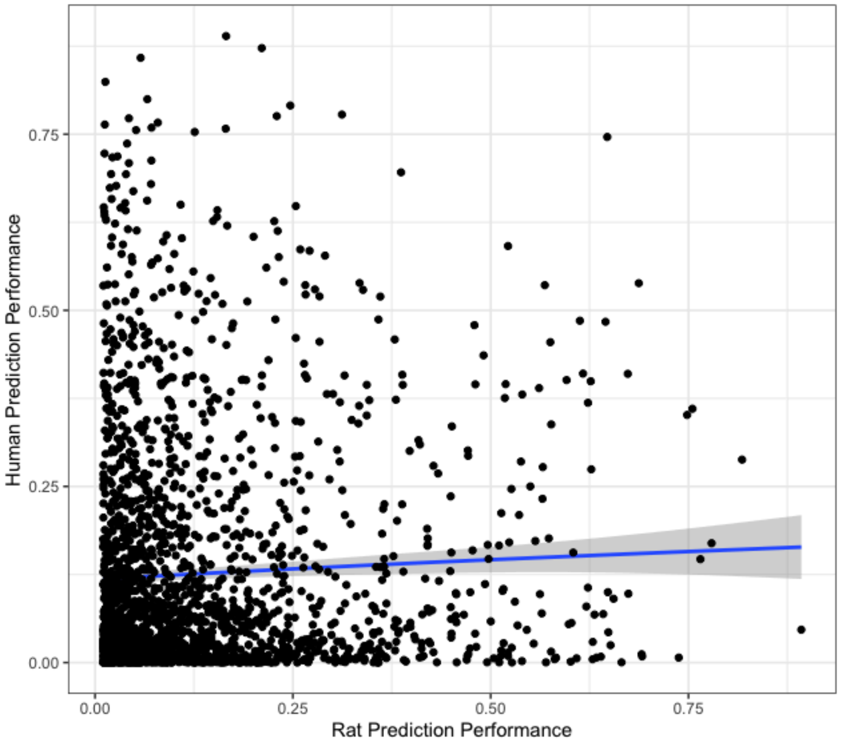


**Figure S3: Shared genetic architecture of gene expression in rats and humans** Prediction performance in humans vs rats. The performance measure (Pearson correlation) was significantly correlated across species ($R$ = 0.06, $P$ = 8.03 x $10^{-6}$).

.

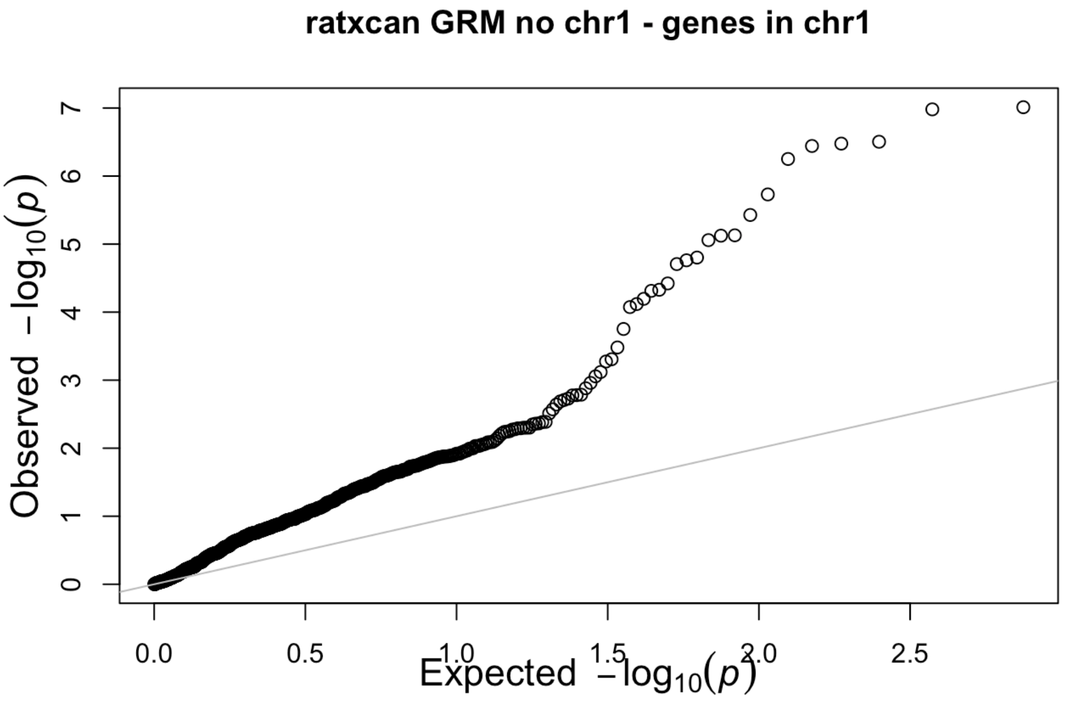


**Figure S4: Leaving one chromosome out for GRM calculation under corrects the inflation.** Using the same null trait simulation used in Figure 3, we performed RatXcan association for genes in chromosome 1 using a GRM calculated excluding variants in chromosome 1. Results for genes in chromosome 1 are shown. A well corrected QQ-plot should have all points on the gray diagonal line but we observed apparent inflation. Hence LOCO for GRM calculation is not recommended.

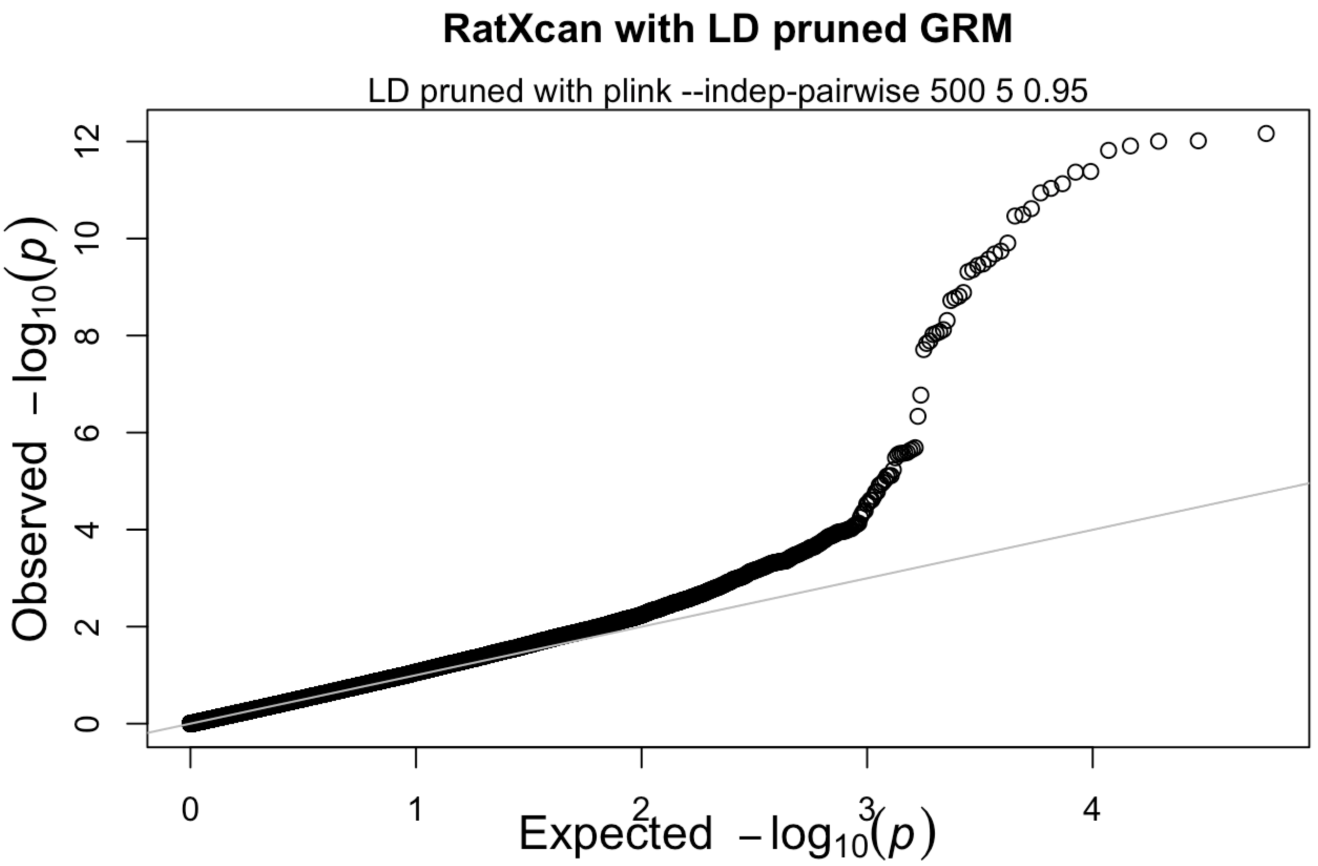


Figure S5: Calculating GRM with LD pruned variants () reduces the effectiveness of the correction. Using the same null trait simulation shown in Figure 3, we performed RatXcan using an LD-pruned GRM. Pruning was done using plink with –indep-pairwise 500 5 0.95, which retains variants with r2 smaller than 0.95 using window size of 500Kb and shifting the window 5 variants at a time. A well corrected QQ-plot should have all points on the gray diagonal line but we observed apparent inflation. Hence LD-pruning is not recommended.

### Supplementary Tables

**Supplementary Table 1**: Body length association with predicted gene expression for the 5 brain regions (Infralimbic Cortex, IL; Lateral Habenula, LHb; Prelimbic Cortex, PL; Orbitofrontal Cortex, OFC).

Column name annotation

gene_name: gene name

p_acat_5: Combined p-values across 5 brain regions using the ACAT method

chr: chromosome

start: start position of the gene

qval: qvalue calculated with the qvalue package

p_human: p-value of the association in humans of the mapped human trait (phenomexcan.org)

hugo_gene: mapped human gene name

trait: rat trait name

gene: rat gene ensembl id

gene_id: mapped human gene ensembl id

AC: p-value of association with predicted expression in Nucleus Accumbens

IL: p-value of association with predicted expression in Infralimbic Cortex

LH: p-value of association with predicted expression in Lateral Habenula

PL: p-value of association with predicted expression in Prelimbic Cortex

OFC:  p-value of association with predicted expression in Orbitofrontal Cortex

**Supplementary Table 2**: Body Mass Index association with predicted gene expression for the 5 brain regions (Infralimbic Cortex, IL; Lateral Habenula, LHb; Prelimbic Cortex, PL; Orbitofrontal Cortex, OFC).

Column name annotation is the same as Supplementary Table 1.
